# Attention in Irritable Bowel Syndrome: A Systematic Review of Affected Domains and Brain-Gut Axis Interactions

**DOI:** 10.1101/2025.01.05.631376

**Authors:** Reyhaneh Akbari, Yeganeh Salimi, Fatemeh Dehghani Aarani, Ehsan Rezayat

## Abstract

**Background:** Irritable Bowel Syndrome (IBS) is a prevalent functional gastrointestinal disorder characterized by abdominal pain and altered bowel habits, significantly impacting patients’ quality of life. Recent research suggests that attention may be affected in individuals with IBS, potentially influencing symptom perception and emotional distress.

**Objective:** This systematic review aims to investigate the relationship between attention and IBS, focusing on the affected domains of attention and the interactions within the brain-gut axis.

**Methods:** A comprehensive literature search was conducted across multiple databases, including MEDLINE/PubMed, PsychINFO, and Scopus, from January 1990 to December 2024. Studies were included if they assessed attention in adult IBS patients and employed valid measurement tools. A total of 24 studies were selected for analysis, encompassing various methodologies, including neuroimaging and behavioral assessments.

**Results:** The findings indicate that IBS patients exhibit significant attentional biases, particularly towards gastrointestinal-related stimuli, reflecting heightened sensitivity and hypervigilance. Specific domains of attention, including selective attention, sustained attention, and pre-attentional processing, were identified as being affected. The review highlights the role of psychological factors, such as anxiety and depression, in modulating attention in IBS. Neuroimaging studies revealed altered brain activation patterns in regions associated with attention and emotional processing, suggesting a complex interplay between cognitive function and the brain-gut axis.

**Conclusion:** This systematic review underscores the multifaceted nature of attention in IBS, revealing specific attentional deficits and biases that may contribute to symptom exacerbation and emotional distress. The findings emphasize the need for further research to explore the underlying mechanisms and potential therapeutic interventions aimed at addressing attention in IBS patients.

## 1. Introduction

Irritable bowel syndrome (IBS) is a chronic functional bowel disorder characterized by changes in stool form or frequency, often accompanied by abdominal discomfort or pain (Lacy et al., 2016). It affects approximately 1.5% to 10% of the global population, depending on the diagnostic criteria used (Oka et al., 2020). The symptoms of IBS significantly impair quality of life, with individuals experiencing considerable challenges in various life domains compared to healthy individuals(Cassar et al., 2020). Additionally, anxiety and depression are prevalent in IBS patients, affecting up to one-third of those diagnosed (Staudacher et al., 2023). Recent research has emphasized the role of the brain-gut axis, which involves the communication between the central nervous system and the enteric nervous system, in the pathophysiology of IBS (Fichna & Storr, 2012). Advanced imaging techniques have revealed abnormal brain responses in IBS patients, suggesting alterations in cognitive profiles (Mayer, Labus, et al., 2015; Rustamov et al., 2020). Emerging studies are investigating the link between IBS and cognitive function, noting that dysregulation of the hypothalamic-pituitary-adrenal (HPA) axis and chronic stress can impair cognitive abilities (Kennedy et al., 2014). Furthermore, mood disorders, chronic pain, and changes in gut microbiota have been associated with cognitive deficits, particularly in attention and memory (Hu et al., 2021; Moloney et al., 2016).

Attentional processes play a significant role in the perception and management of IBS symptoms. BS patients often display a heightened attentional bias towards pain-related stimuli. This bias is characterized by faster engagement with pain words and increased focus on gastrointestinal symptoms, which can lead to a vicious cycle of symptom perception and illness behavior (Gibbs-Gallagher et al., 2001). Such attentional biases are linked to increased reporting of somatic symptoms and greater illness-related behaviors, such as taking sick leave (Chapman & Martin, 2011).

Neural circuitry dysfunctions in IBS include hyperactivity in the salience network (anterior insula and anterior mid-cingulate cortex) (Liu et al., 2016) and emotional arousal network (pregenual anterior cingulate cortex, amygdala) (Labus et al., 2009; Tillisch et al., 2012), combined with hypoactivity in the central executive network (dorsolateral prefrontal cortex) (Chen et al., 2021; Naliboff et al., 2008). The hypoactivity of the dorsolateral prefrontal cortex is associated with attentional bias and executive dysfunction, while hyperactivity in the salience network influences attention through its connections with emotional and autonomic networks, contributing to visceral hypersensitivity and a heightened stress response (Mayer, Labus, et al., 2015). These disruptions may underlie the attention difficulties experienced by individuals with IBS.

Despite the growing body of literature on the cognitive aspects of IBS, there has been a lack of comprehensive reviews specifically focusing on attention and its affected domains in this population. Previous studies have shown that IBS patients may experience impairments in various cognitive functions, including selective attention, sustained attention, and hypervigilance. However, the specific nature and extent of these cognitive deficits remain poorly understood. Moreover, two systematic reviews have recently addressed cognitive aspects of IBS, providing valuable insights into the relationship between cognitive function and IBS symptoms (Lam et al., 2019; Wong et al., 2019). However, both reviews call for further investigation into the specific cognitive mechanisms and their implications for symptom management. Despite these contributions, a significant gap exists in the literature regarding a comprehensive examination of attention and its various aspects in IBS. While previous reviews have identified attentional biases, they have not systematically synthesized findings across different attention domains or explored how these biases interact with the brain-gut axis. This lack of focused research limits the understanding of how attention influences symptom perception and management in IBS patients

This systematic review aims to address this gap by providing a thorough overview of the existing research on cognitive attention in IBS patients. By synthesizing findings from multiple studies, this review will elucidate the cognitive mechanisms underlying IBS. Understanding these cognitive processes is essential for developing targeted interventions that address both the psychological and physiological challenges faced by individuals with IBS, ultimately improving their quality of life.

## 2. Methods

This systematic review was conducted following the PRISMA guidelines. The protocol was registered in PROSPERO under the Registration Number CRD42024584371 focusing on irritable bowel syndrome (IBS), attention, its domains and possible brain-gut interactions.

### 2.1. Search Strategy

Relevant published research was identified through two independent searches of the following electronic databases: MEDLINE/PubMed, PsychINFO, and Scopus from 1 January 1990 to December 2024 (Page et al., 2021). Relevant dissertations, along with secondary and conference papers from Scopus, were also reviewed as sources of gray literature. We updated our search twice—initially in July 2024, and subsequently on December, 2024—to encompass studies published in PubMed up until December 2024. Additionally, we manually reviewed reference lists from relevant reviews and meta-analyses

The following search terms were used in all of the mentioned databases:(“Irritable Bowel Syndrome” OR “IBS” OR “Functional Gastrointestinal Disorder” OR “FGID”) AND (“Cognitive Function” OR “Cognitive Impairment” OR “Attention” OR “Attention Bias” OR “Selective Attention” OR “Sustain Attention” OR “Executive Function” OR “Cognitive Processes” OR “Attention Network” OR “Brain-Gut Axis” OR “Gut-Microbiota-Brain Axis”)Supplementary Table1 shows results for each string. 911 articles were retrieved. Titles, abstracts, and full texts of articles were screened independently by two reviewers (R.A. and Y.S)

### 2.2. Selection Process

The following categories were excluded: news, books, reviews, editorials, case studies, and literature reviews. After removing duplicates, 210 articles remained, which were independently screened by 2 authors (R.A. and Y.S.) by reading abstracts and full text. Disagreement was resolved by discussion between the 2 independent authors; if no agreement was reached, a third independent party (E.R.) was involved as an arbiter. Twenty-four studies were selected (Fig. 1). Results are presented divided up by methodology in Table 1.

**Figure 1.**
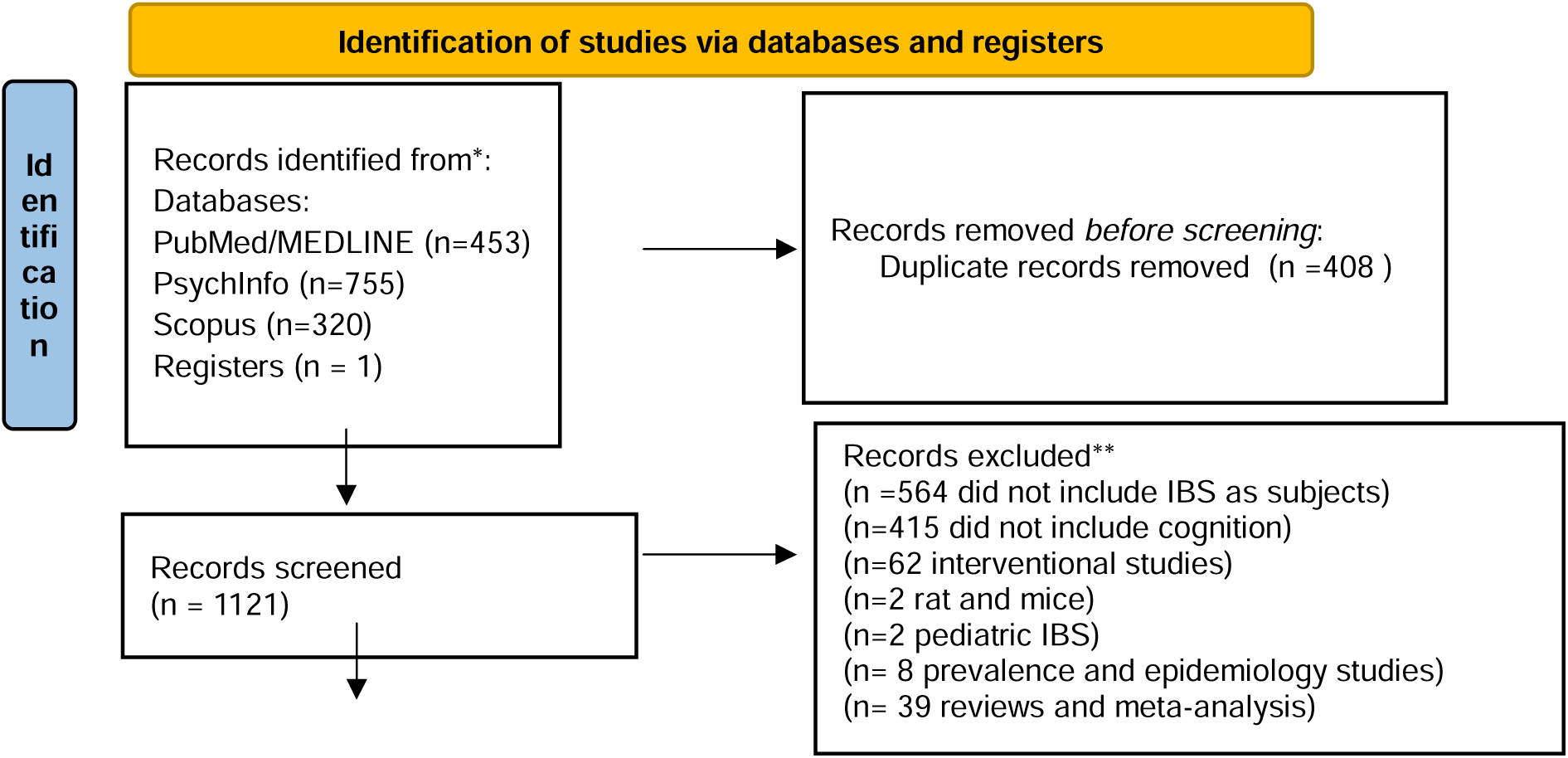
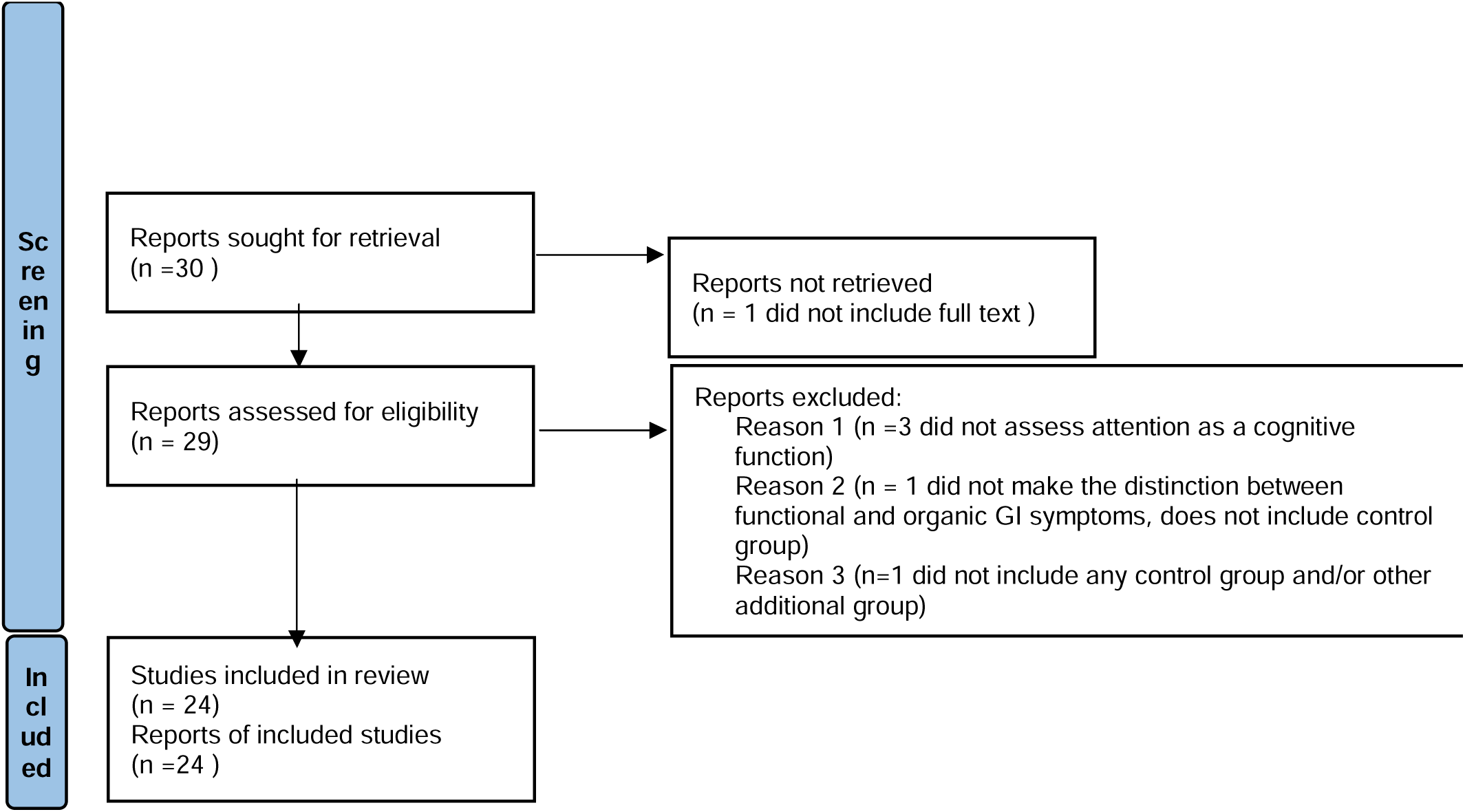
Preferred Reporting Items for Systematic Reviews and Meta-analyses (PRISMA) flow diagram illustrating the bibliographic search and the selection process.

**Table 1:**
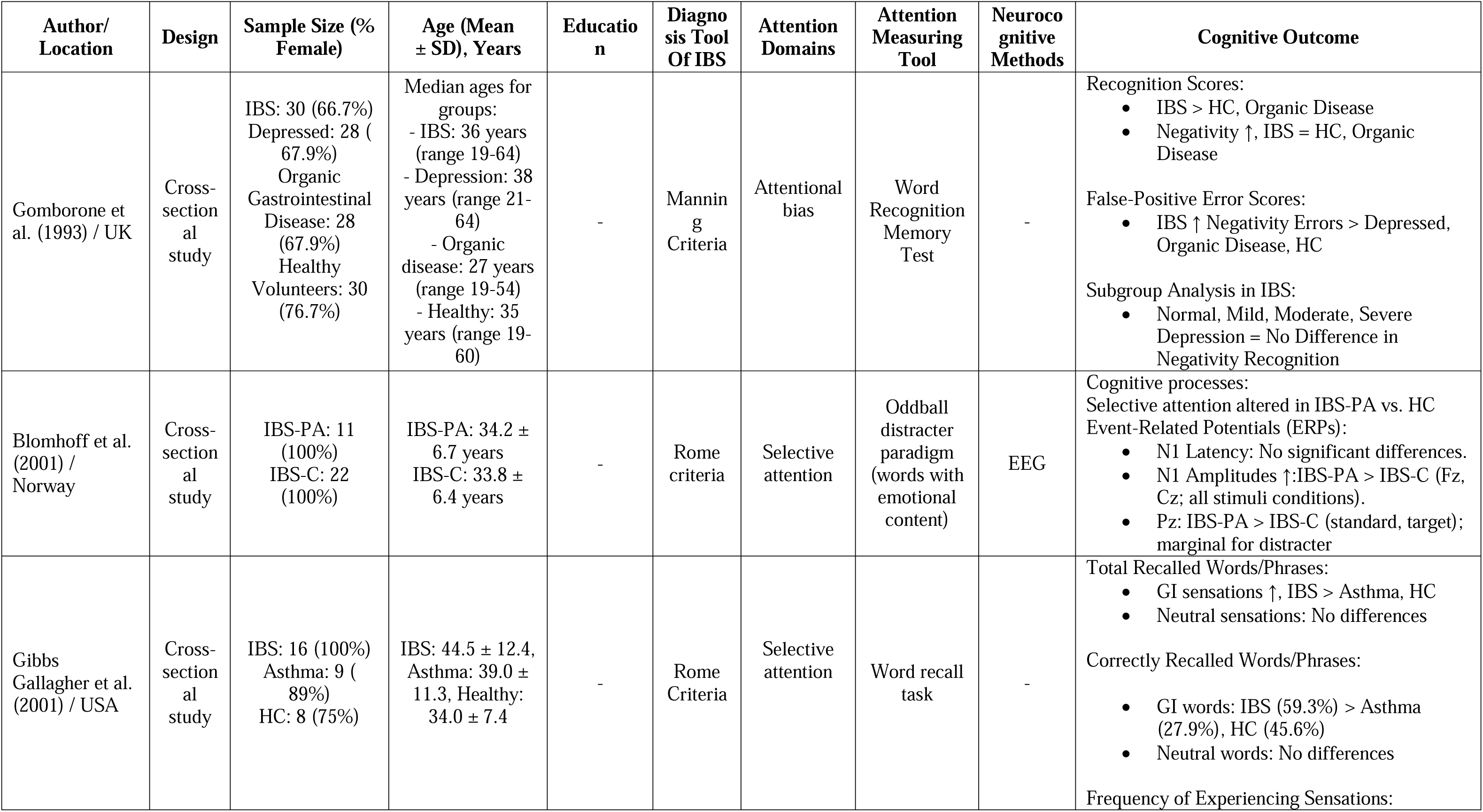
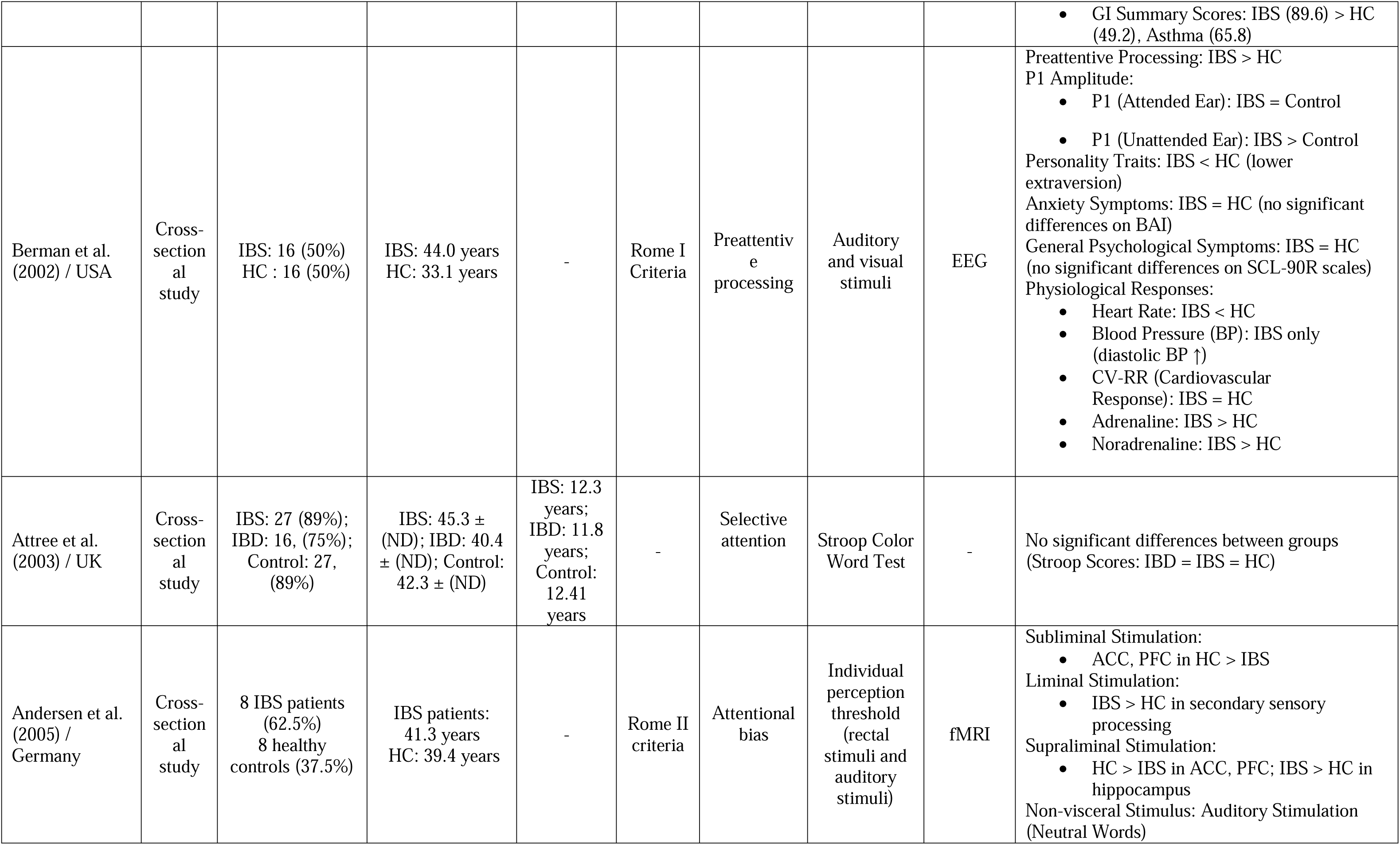
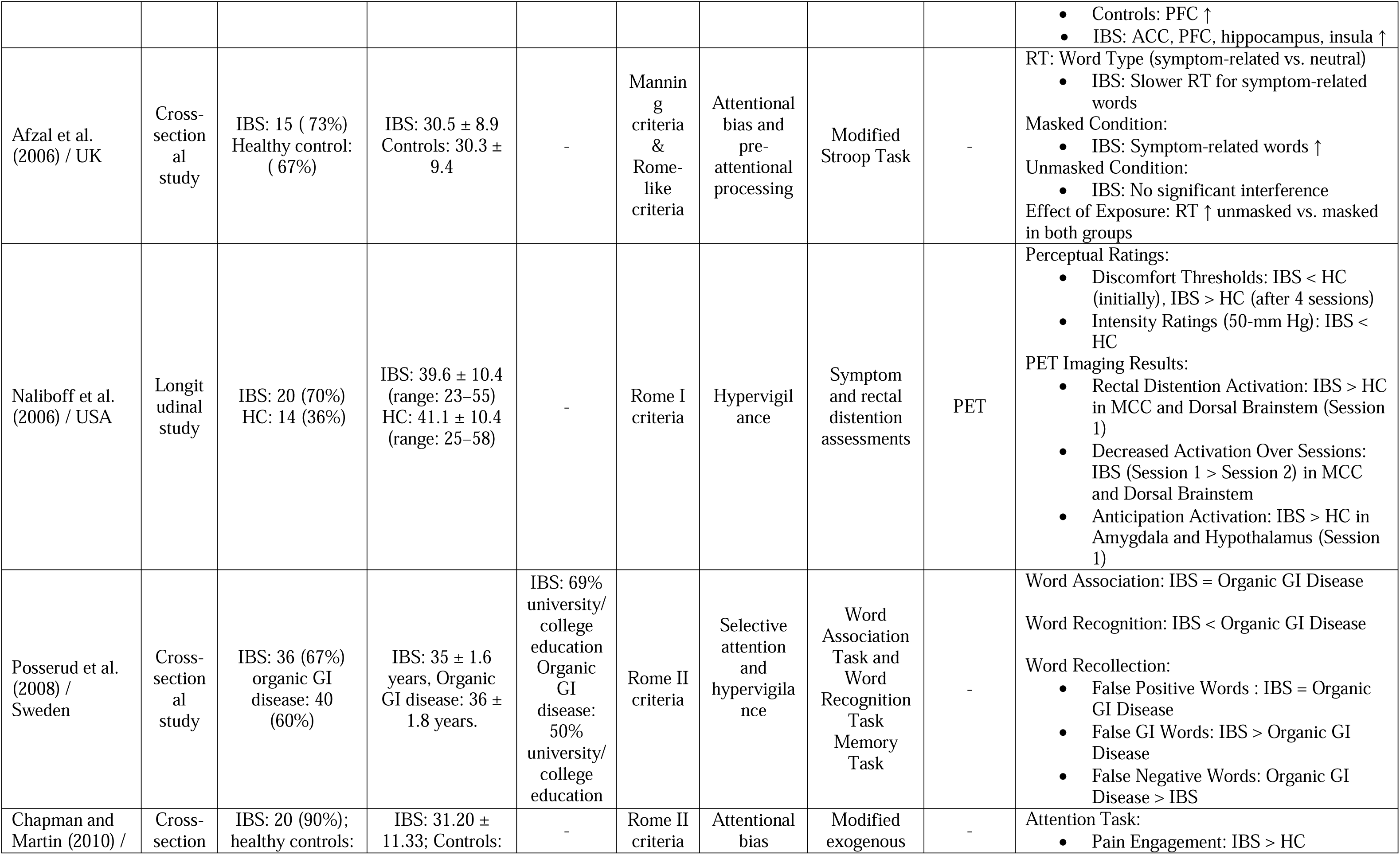
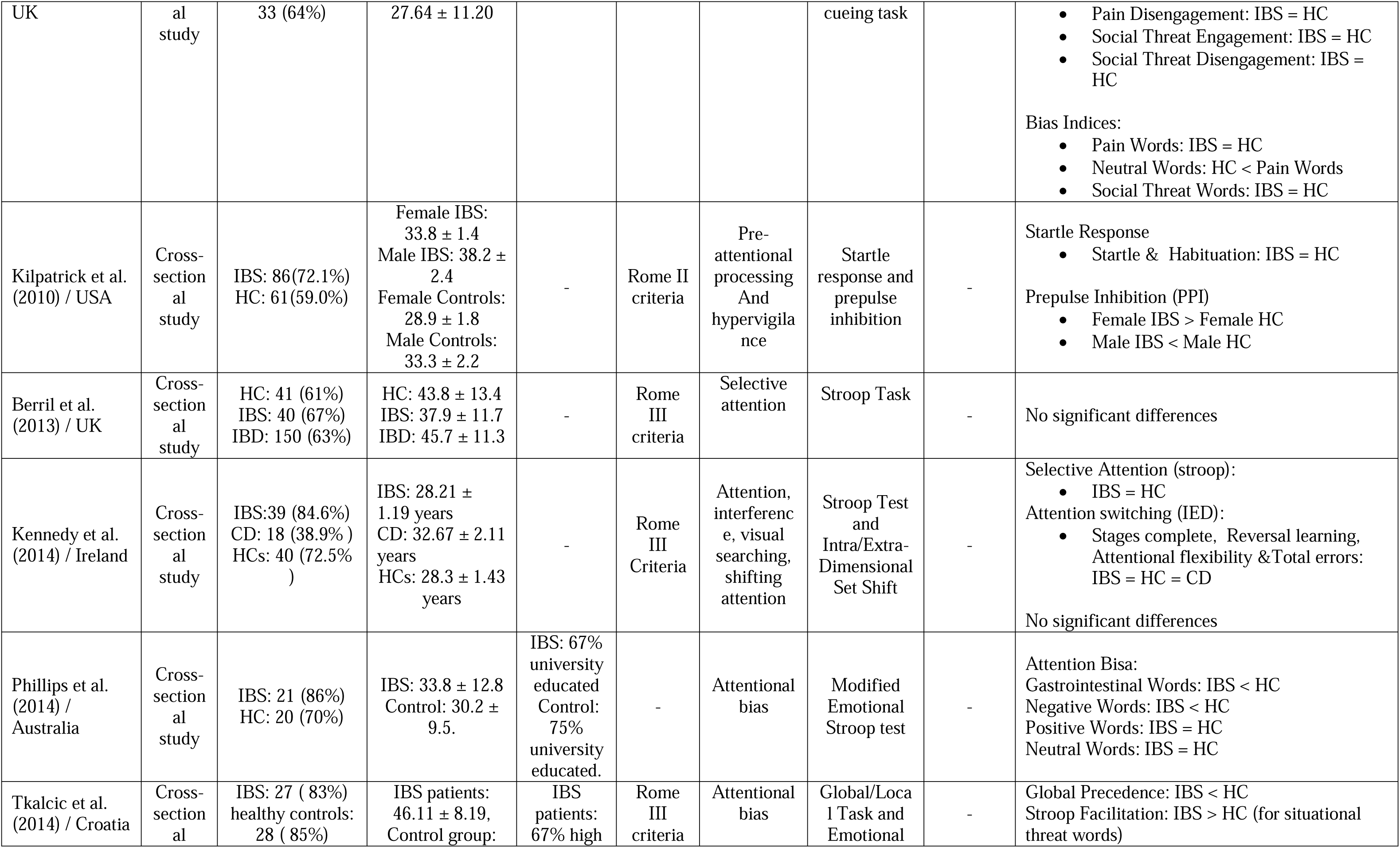
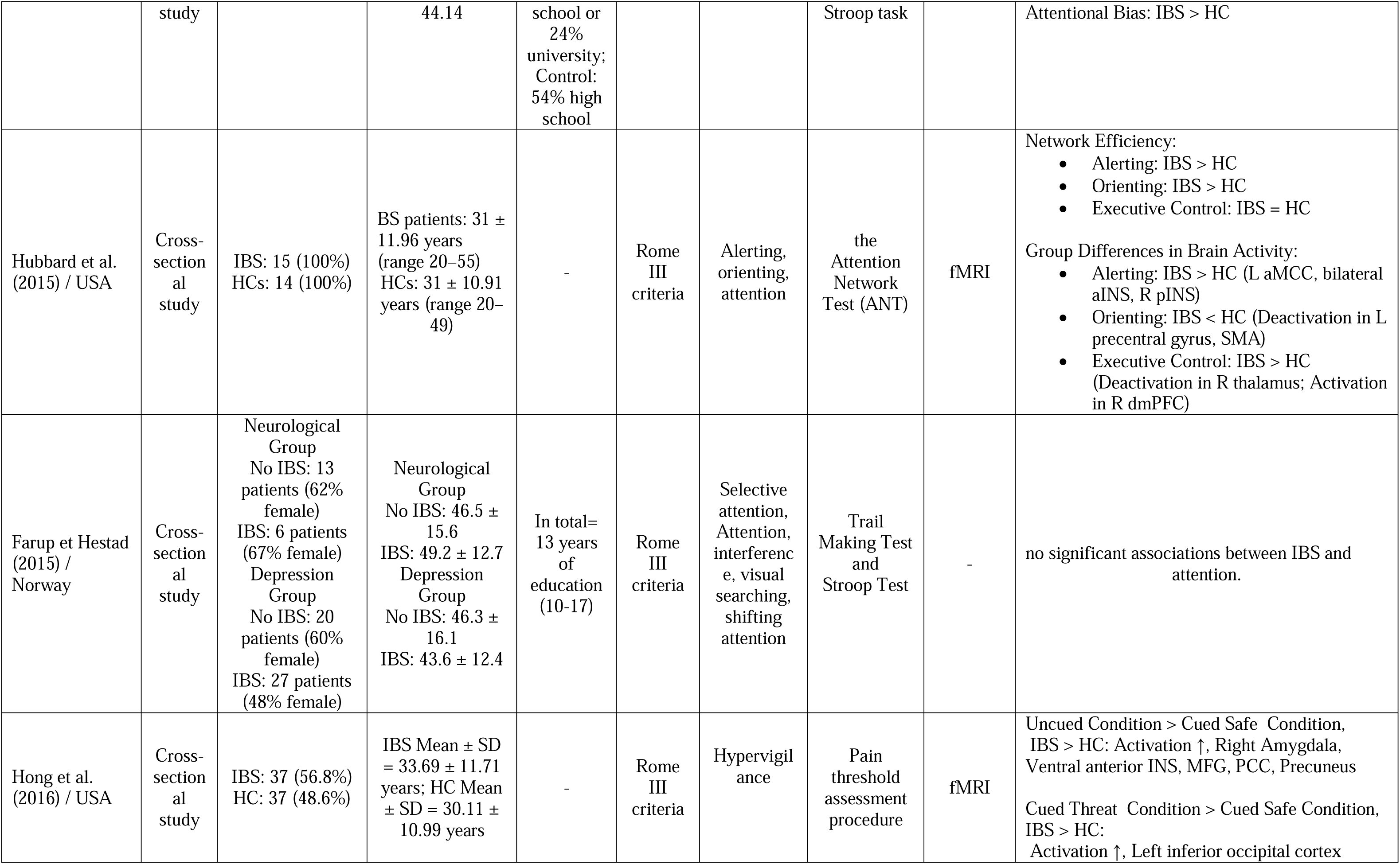
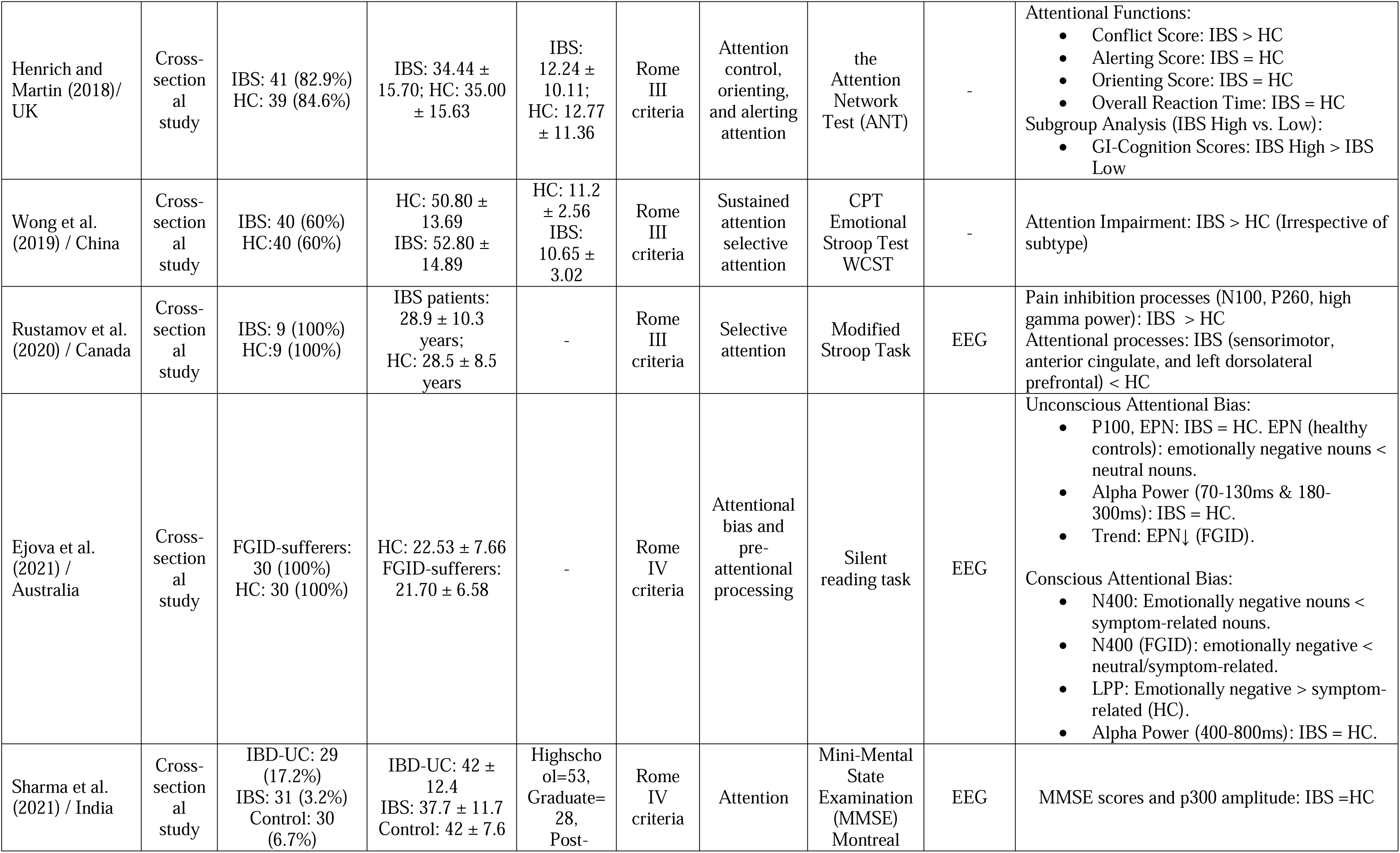
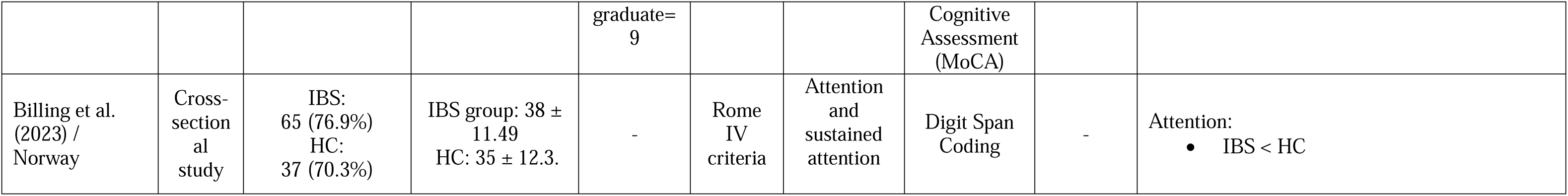
Characteristics and Key Findings of Studies on Cognitive Attention in IBS.

Inclusion criteria were: (1) Full text English-written articles, (2) articles including only human subjects above 18 years old, (3) studies on IBS, and (4) studies that assessed attention as a cognitive function. Exclusion criteria were: (1) interventional in design, (2) studies solely about any other functional somatic disorder, (3) review articles, (4), studies without comparison group, and (5) studies including pediatric patients or non-human subjects with IBS. The search was conducted using the following keywords: (irritable bowel syndrome), (IBS), (functional gastrointestinal disorders), and (nervous colon); and combined on a string with the keywords: (attention), (selective attention), (attentional bias) and (executive function). Supplementary Table shows results for each string. 911 articles were retrieved. Titles, abstracts, and full texts of articles were screened independently.by two reviewers (R.A. and Y.S)

### 2.3. Data Collection Process

Data were extracted from selected studies using an Excel sheet created by Y.S., and then verified by R.A. In cases of disagreement, the second author (E.R.) was consulted. To comprehensively evaluate the existing literature on IBS, a range of study characteristics was examined. These included the population studied, study design, IBS subtypes, comparison groups, sample size, demographics (sex, age, country and education), medications used, attention domains assessed, diagnostic criteria, IBS type, cognitive measuring tools, psychological assessment methods, and neurocognitive techniques and the reported outcomes. By considering these factors, present study aimed to provide a thorough overview of the available evidence on IBS and its association with cognitive attention, psychological, and neurocognitive factors.

### 2.4. Eligibility Criteria

Inclusion criteria were: (1) Full text English-written articles, (2) articles including only human subjects above 18 years old, (3) studies on IBS, and (4) studies that assessed attention as a cognitive function. Exclusion criteria were: (1) interventional in design, (2) studies solely about any other functional somatic disorder, (3) review articles, case reports, case series (4), studies without comparison group, (5) studies including pediatric patients or non-human subjects with IBS.

### 2.5. Risk of Bias Assessment and Quality of Evidence

The Joanna Briggs Institute (JBI) Critical Appraisal Checklist for Analytical Cross-Sectional Studies and JBI Critical Appraisal Checklist for Cohort Studies were applied because all eligible studies included in this review had a cross-sectional or longitudinal design (Moola et al., 2015). The complete checklist can be found in Appendix 1. Two reviewers (R.A. and Y.S.) independently assessed the quality of the included studies. Any discrepancies were resolved by the third reviewer (E.R.). Tables 2 and 3 and provide an assessment of the quality of the studies. The average assessment of the quality of cross-sectional studies was 8 points (on a scale of 8 points), and for the cohort studies, it was 11 points (on a scale of 11 points). The questions were answered with the options ’Yes’, ’No’, ’Unclear’, or ’Not applicable’. A score of one was given for each ’Yes’ response, while a score of zero was assigned for ’No’ or ’Unclear’ responses. The maximum possible score was 8 for cross-sectional studies and 11 for cohort studies. Overall scores for each paper were calculated as percentages, and study quality was rated as high (80-100%), fair (50-79%), or low (<50%) (Moola et al., 2015). Of the 24 included studies, 20 were rated as high quality, while 4 were rated as fair quality. None of the studies were rated as low quality nor were rejected based on quality assessments.

**Table 2:**
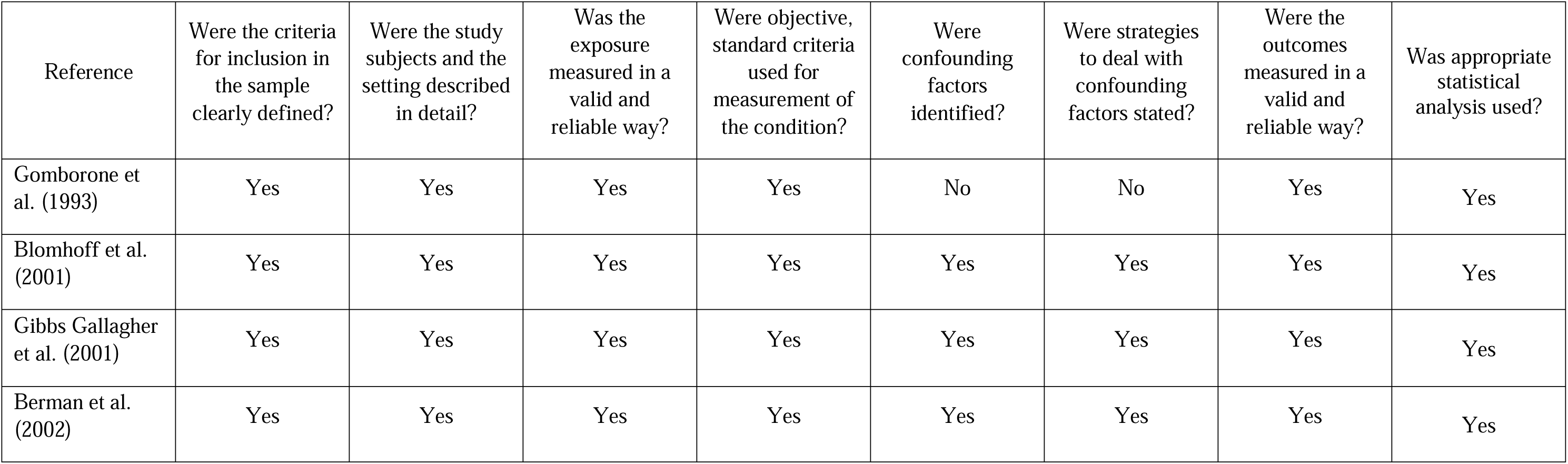
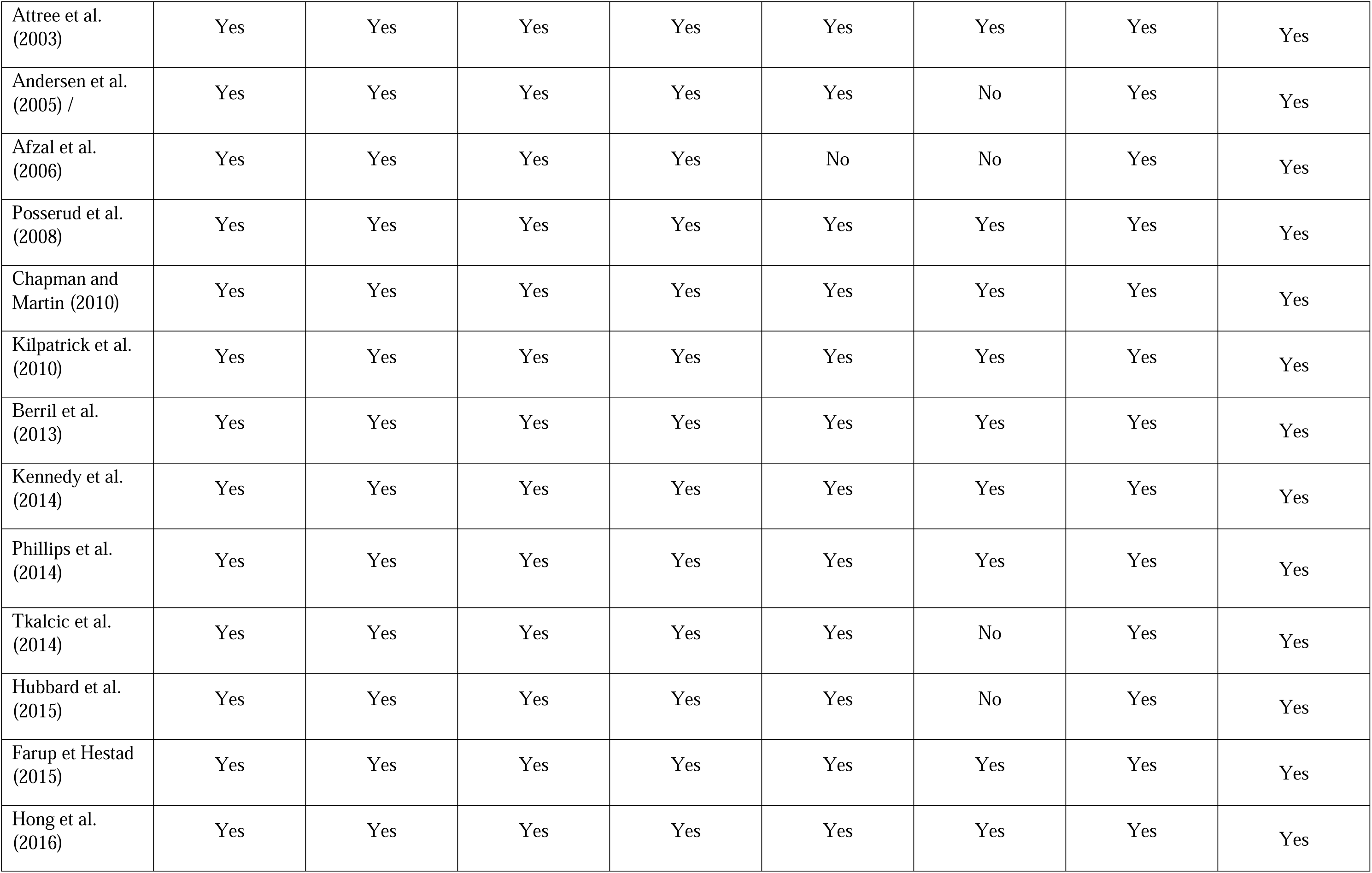
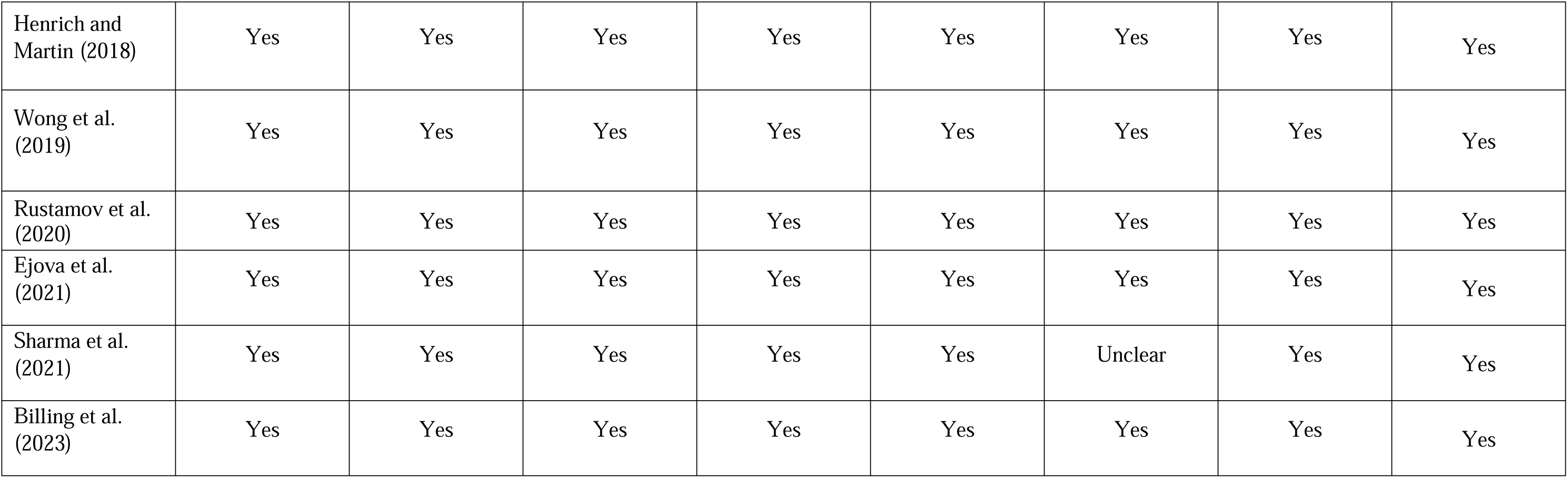
Summary of the JBI Critical Appraisal Checklist for Analytical Cross-Sectional Studies.

**Table 3:** Summary of the JBI Critical Appraisal Checklist for Cohort Studies.

| References | Were the two groups similar and recruited from the same population? | Were the exposures measured similarly to assign people to both exposed and unexposed groups? | Was the exposure measured in a valid and reliable way? | 4. Were confounding factors identified? | 5. Were strategies to deal with confounding factors stated? | Were the groups/participants free of the outcome at the start of the study (or at the moment of exposure)? | Were the outcomes measured in a valid and reliable way? | Was the follow up time reported and sufficient to be long enough for outcomes to occur? | Was follow up complete, and if not, were the reasons for loss to follow up described and explored? | 10. Were strategies to address incomplete follow up utilized? | 11. Was appropriate statistical analysis used? |
| --- | --- | --- | --- | --- | --- | --- | --- | --- | --- | --- | --- |
| Naliboff et al. (2006) / USA | Yes | Yes | Yes | Yes | No | Yes | Yes | Yes | No | No | Yes |

## 3. Results

Due to the variation in study designs and outcome assessments, the results are presented narratively, supplemented by Table 1.

### 3.1. Search Result

Figure 1 shows our search result and screening results according to PRISMA. Our search identified 1529 records from databases and registers. After removing 408 duplicates, 1121 records were screened by title and abstract. Following this review process, 30 reports were sought for retrieval, but one report was not retrieved as it did not include the full text. Thus, 29 reports were assessed for eligibility. Of these, 24 studies were included in the review. The excluded publications did not fulfill the inclusion criteria for various reasons, such as not assessing attention as a cognitive function, not distinguishing between functional and organic GI symptoms, or not including a control group. We concluded that 24 papers could be included in our systematic review. The details are shown in the PRISMA flow diagram of Figure 1.

The countries in which the research work of the 24 selected articles was carried out show a high degree of heterogeneity. The major origins were the United Kingdom (29%) and the United States (29%). These were followed by Norway (13%) and Australia (8%). Other countries included Germany, Sweden, Croatia, Ireland, China, Canada, and India (4% each).

### 3.2. Study Characteristics

The majority of studies were cross-sectional (23), with only one adopting a longitudinal design. The total number of participants evaluated across the 24 studies is 1,576. Across these studies, the number of healthy controls (HC) varies from 8 to 61, while the number of participants with irritable bowel syndrome (IBS) ranges from 8 to 86. Several studies also include participants with other conditions such as depression (Gomborone et al., 1993), anxiety (Blomhoff et al., 2001), asthma (Gibbs-Gallagher et al., 2001), and organic gastrointestinal diseases like Crohn’s disease (Kennedy et al., 2014) and ulcerative colitis (Attree et al., 2003; Gomborone et al., 1993; Posserud et al., 2009).

Diagnosis of IBS was primarily based on the Rome criteria (I–IV) across most studies, though a few employed the Manning criteria or similar diagnostic tools to confirm IBS. In 19 out of the 24 articles (79%), a health professional diagnosed IBS alongside criteria like Rome, IBS-SSS, and Manning. In the remaining 5 articles (Billing et al., 2024; Ejova et al., 2021; Farup & Hestad, 2015; Posserud et al., 2009; Tkalcic et al., 2014), while the research was conducted within a healthcare setting, the clinical criteria for IBS diagnosis were not specified by a clinician. A variety of tools were used to measure attention, 10 studies used the Stroop Task or Modified Stroop Task 3 Word Recognition Memory Task 3 individual perception threshold (symptom and rectal distention assessments), 2 Attention Network Tests (ANT), 2 oddball distracter paradigm others used Modified exogenous cueing task (Chapman & Martin, 2011), the Trail Making Test (TMT), Continuous Performance Test (CPT), Wisconsin Card Sorting Test (WCST), silent reading, Digit Span Coding, The Global/Local Task, Mini-Mental State Examination (MMSE), Montreal Cognitive Assessment (MOCA) auditory and visual stimuli, and startle response and prepulse inhibition.

All the included studies consisted of 409 participants diagnosed with IBS. The proportion of female participants ranged from 50% to 100% of subjects. The mean age of IBS participants ranged from 21.7 to 45.3 years, with a standard deviation of 12.8 years. The mean education level ranged from 10.27 to 14.59 years, and 15 studies did not provide information on level of education. The control group consisted of participants with a mean age of 30.2 years and a standard deviation of 9.5 years.

### 3.3. Behavioral analyses of Attention Domains

The studies included in the review varied in methodology, patient characteristics, outcome measures, and the tools used to assess these outcomes. For instance, four different research groups each used a unique test to evaluate attention (as shown in Table 1). This diversity created challenges for data synthesis in a meta-analysis. Additionally, for certain cognitive domains, only two or three studies were available, often resulting in inconsistent findings.

Due to this heterogeneity in metrics, outcomes, and participant profiles, we opted not to conduct a meta-analysis. Instead, we performed a narrative analysis, organizing the findings by attention domain. Here, we assessed the IBS attentional deficits in different attentional domain separately.

#### 3.3.1 Attention

Two studies assessed attention using Digit Span Coding, Mini-Mental State Examination, Montreal Cognitive Assessment (Billing et al., 2024; Sharma et al., 2021). These studies did not establish a strong or direct association between IBS and attention deficits. While IBS patients may show slight reductions in attention performance, as indicated in the Billing et al (2024), these differences are not statistically significant and could reflect subtle or secondary effects rather than a robust impact on attention itself. Depression may play an indirect role in influencing attention in some IBS patients, as suggested by Farup and Hestad (2015), but this seems to be influenced by the specific nature of depression associated with IBS.

Comparatively, attention-related impairments are more evident in IBD patients, which was not included in our systematic review, where cognitive processing speed and general cognitive scores are lower, suggesting that cognitive challenges may be more characteristic of IBD than IBS. Thus, while IBS may occasionally correlate with mild cognitive effects, including attention, these effects are inconsistent and generally less pronounced than those observed in other gastrointestinal disorders like IBD (Sharma et al., 2021).

#### 3.3.2. Selective Attention

Selective attention restricts neural processing and behavioral responses to a relevant subset of available stimuli, while excluding other valid stimuli from consideration (Krauzlis et al., 2018). Eight studies examined selective attention measuring with Modified Stroop Task, oddball distracter paradigm, Word Recognition Memory Task and Stroop task (Attree et al., 2003; Blomhoff et al., 2001; Gibbs-Gallagher et al., 2001).

The collective evidence on selective attention in IBS presents a nuanced picture, with studies showing both intact general attention processes and specific attentional biases. While broad cognitive impairments are generally absent in IBS patients (Attree et al., 2003; Berrill et al., 2013; Farup & Hestad, 2015), there is evidence of selective attentional biases towards GI-related stimuli (Gibbs-Gallagher et al., 2001; Kennedy et al., 2014; Posserud et al., 2009). This selective focus on symptom-related cues aligns with cognitive-behavioral models of IBS, suggesting that patients may have a predisposition to attend more to bodily sensations and GI symptoms, which could reinforce symptom perception and anxiety.

Moreover, while Rustamov et al. (2020) demonstrate that selective attention may not effectively inhibit pain-related brain activity in IBS patients, indicating altered attentional processing in response to pain, Wong et al. (2019) found no specific attentional biases in an emotional Stroop task. This discrepancy may reflect variations in how attention operates in response to symptom-specific stimuli versus general emotional stimuli.

Selective attention in IBS appears to be intact in a general sense, but symptom-related attentional biases are often present. These biases may interfere with patients’ ability to effectively manage pain and other symptoms by directing more attentional resources towards GI cues, potentially intensifying symptom perception and exacerbating the cognitive and emotional challenges associated with IBS.

#### 3.3.3 Attention Bias

Attentional bias is the tendency for patients to pay attention selectively to information related to their current concerns (Crombez et al., 2015) and it is characterized by attending to emotional cues compared to neutral cues, influenced by underlying affect (Howarth et al., 2021). Seven of the reviewed studies assessed attention bias by Word Recognition Memory Task, individual perception threshold, Modified Stroop Task, modified exogenous cueing task, Global/local task and silent reading task (Afzal et al., 2006; Andresen et al., 2005; Chapman & Martin, 2011; Ejova et al., 2021; Gomborone et al., 1993; Phillips et al., 2014; Tkalcic et al., 2014)

The findings across these studies converge on a consistent pattern of attentional biases in IBS patients, emphasizing heightened sensitivity to negative and symptom-related stimuli. IBS patients demonstrate a confirmatory bias towards emotionally negative and pain-related information, often engaging more quickly and making more false-positive errors for negative content compared to healthy controls, as observed in both memory and Stroop tasks (Chapman & Martin, 2011; Gomborone et al., 1993; Phillips et al., 2014). Increased neural activation in emotional processing areas also supports this tendency, suggesting that IBS patients have heightened reactivity to even neutral stimuli, which may generalize to non-gastrointestinal emotional content (Andresen et al., 2005). Additionally, traits like neuroticism, trait anxiety, and visceral anxiety correlate with attentional biases in IBS, especially towards situational threats, indicating a unique vulnerability compared to the general population (Tkalcic et al., 2014). Selective attentional processing appears to operate differently depending on awareness: IBS patients exhibit significant subliminal processing of symptom-related words, but not in conscious tasks (Afzal et al., 2006) or these biases may manifest at both unconscious and conscious levels, as reflected in heightened occipital and N400 amplitudes, which correlate with psychosocial factors like anxiety and depression (Ejova et al., 2021).

Together, these findings suggest that attentional biases in IBS and FGID patients are multifaceted, potentially reinforcing symptom perception, emotional distress, and maladaptive illness behaviors through both automatic and conscious processing pathways.

### 3.3.4. Sustain Attention

Sustained attention is the process of maintaining response persistence and continuous effort over extended periods of time (Ko et al., 2017). Sustain attention is only assessed in two if the included studies assessing with Continuous Performance Test and Digit Span Coding (Billing et al., 2024; Wong et al., 2019). The evidence on sustained attention in IBS patients suggests that while broad cognitive impairments are generally minimal, there are specific, stable deficits in sustained attention that may be a core feature of the condition. Wong et al. (2019) found that IBS patients exhibited increased variability in reaction times (SDRT) on the Continuous Performance Test, indicating a significant impairment in sustained attention compared to healthy controls. This attentional difficulty was consistent over time and did not correlate with the severity or chronicity of IBS symptoms, suggesting it may be a trait characteristic of IBS rather than directly influenced by symptom fluctuations. Billing et al. (2023) further supports these findings by revealing that IBS patients have lower scores on cognitive domains such as immediate memory and recall, along with high levels of anxiety and depression. However, there was no significant correlation between emotional symptom severity and cognitive performance in this group, implying that attentional deficits and emotional symptoms may represent independent domains of dysfunction. Together, these findings suggest that while IBS patients may not experience global cognitive deficits, impairments in sustained attention could be stable, trait-like characteristics that warrant attention in clinical assessments.

#### 3.3.5. Hypervigilance

Hypervigilance is a state of heightened awareness and watchfulness, associated with exposure threat (Smith et al., 2019) which can involve in enhanced or exaggerated search of environmental stimuli or scan for threatening information (Rollman, 2009). Four studies investigated hypervigilance. Startle response and prepulse inhibition, individual perception threshold and Word Recognition Memory Task was used (Hong et al., 2016; Kilpatrick et al., 2010; Naliboff et al., 2008; Posserud et al., 2009). The evidence on hypervigilance in IBS reveals a pronounced, multifaceted attentional sensitivity toward gastrointestinal (GI) sensations and potential threats, underscoring a complex interaction between sensory processing and cognitive evaluation. Posserud et al. (2009) demonstrate that IBS patients display faster recognition of GI-related and negatively valenced words compared to patients with organic GI diseases, as well as a tendency to recall more false GI words, highlighting their selective attentiveness toward GI and emotionally charged information. This heightened vigilance appears to be generalized, as IBS patients also quickly recognize positive emotional and non-GI-related stimuli, suggesting an elevated baseline of attentional sensitivity. Kilpatrick et al. (2010) extend this understanding by revealing that IBS patients, particularly females, show enhanced sensorimotor gating (PPI) responses to visceral sensations, indicating an intensified state of alertness and hyper-attention to potential threats in the environment.

Hong et al. (2016) further corroborate this heightened salience detection, showing greater activation in the Middle Frontal Gyrus and other attentional processing regions during both anticipated and unanticipated threat cues. This activation pattern reflects increased engagement in evaluating potential threats, particularly pain-related, as a key component of hypervigilance in IBS. Finally, Naliboff et al. (2008) add a longitudinal perspective by demonstrating that while perceptual responses to visceral threats can normalize with repeated exposure, central arousal networks (including limbic and pontine regions) show decreased activity over time. This suggests that, despite an initial heightened response to visceral threats, IBS patients may eventually habituate to such stimuli, with selective regions stabilizing in activation but maintaining sensitivity to threat anticipation.

Together, these studies indicate that hypervigilance in IBS is characterized by an initial heightened sensory and cognitive response to GI sensations and potential threats, with strong activation in threat-related brain networks that may persist over time. This hypervigilance may be underpinned by beliefs about illness severity and an exaggerated sense of threat, which fuel anxiety and symptom-focused attention.

#### 3.3.6. Attention Control, Orienting, and Alerting Attention

Alerting attention maintains an alert state and responds to warning signals; orienting selects information among sensory inputs; and attention control, resolves conflicts and controls thoughts or behaviors (Spagna et al., 2015) and was investigated in four studies (Farup & Hestad, 2015; Henrich & Martin, 2018; Hubbard et al., 2015; Kennedy et al., 2014)

The studies collectively illustrate a pattern of hypervigilance and attentional bias in IBS, with specific attentional sensitivities emerging against a background of mostly intact cognitive flexibility. Kennedy et al. (2014) by using Intra/Extra-Dimensional Set Shift of CANTAB observed that IBS patients showed slight impairments in cognitive flexibility, particularly in attentional set-shifting, suggesting they may have difficulty adjusting focus and disengaging from symptom-related or threat cues. This finding aligns with the tendency for IBS patients to sustain attention on bodily symptoms, potentially intensifying hypervigilance.

At the same time, Hubbard et al. (2015) using the Attention Network Test (ANT) found that IBS patients demonstrated heightened efficiency in orienting and alerting attention networks, indicating they respond more quickly and remain more alert to potential cues in their environment than controls, which supports the concept of a heightened sensitivity to sensory inputs. Contrastingly, Farup and Hestad (2015) with Trail Making Test reported no significant cognitive flexibility deficits on a set-shifting task, suggesting that any attentional bias in IBS patients does not extend to generalized cognitive inflexibility. Similarly, by using ANT, Henrich and Martin (2018) identified lower attentional control in IBS patients but found no differences in orienting or alerting attention compared to healthy participants.

Overall, these findings suggest that while general cognitive functions like set-shifting may not be broadly affected, IBS patients display a selective attentional hypervigilance, especially in orienting and alerting to stimuli. This hypervigilance could reflect a sensitivity to symptom or threat-related cues, reinforcing a cycle of symptom-focused attention and anxiety in IBS.

#### 3.3.7. Pre-Attentional Processing

Pre-attentive processing is the subconscious accumulation of information from the environment (Wolfe & Utochkin, 2019). Pre-attentional processing was assessed in four studies (Afzal et al., 2006; Berman et al., 2002; Ejova et al., 2021; Kilpatrick et al., 2010).

In examining the role of pre-attentional processing in IBS, findings across studies reveal a distinct pattern of heightened, unconscious sensitivity to both sensory and GI-related cues in IBS patients compared to healthy controls. Afzal et al. (2006) using Modified Stroop Task found that IBS patients exhibit an automatic attentional bias towards GI-related words in subliminal (masked) conditions, implying an unconscious attunement to symptom-related stimuli that is activated pre-attentively. Unlike in the supraliminal (unmasked) condition, where conscious processing allowed patients to override this bias, the masked condition suggests that IBS patients have difficulty inhibiting attentional responses to GI cues when they are processed at an unconscious level. This pattern contrasts with healthy controls, who displayed an attentional bias only in the unmasked condition, indicating that IBS patients uniquely exhibit pre-attentional bias towards symptom-related information.

Kilpatrick et al. (2010) examined pre-attentional processing with through prepulse inhibition (PPI), finding notable sex differences. Female IBS patients showed greater PPI than female controls, reflecting heightened pre-attentional processing and possibly hypervigilance to irrelevant stimuli. Male IBS patients, however, demonstrated reduced PPI, indicating difficulty in inhibiting responses to stimuli, which may signal a deficit in pre-attentional filtering. This sex-specific variation was further influenced by symptom severity, which correlated positively with PPI in female patients but negatively in males, suggesting that symptom severity modulates the degree of pre-attentional sensitivity in IBS. The study also noted that menstrual cycle status affected PPI in females, with naturally cycling patients showing enhanced PPI, reinforcing the complexity of sex-related factors in pre-attentional processing in IBS.Berman et al. (2002) highlighted heightened pre-attentional sensitivity in IBS patients via increased P1 amplitudes in response to auditory stimuli, across both attended and unattended conditions. This consistent increase in P1 amplitude across conditions suggests that IBS patients experience an overall elevation in central nervous system reactivity, which aligns with the characteristics of hypervigilance and reduced sensory gating. This lack of sensory gating points to a diminished ability to filter out repetitive or irrelevant stimuli, leading to enhanced processing of all incoming sensory information. Berman’s findings underscore that IBS patients have an augmented, automatic response to sensory input that is pervasive, regardless of task demands, indicating an underlying heightened sensitivity in their pre-attentional systems.Similarly, Ejova et al. (2021) examined pre-attentional processing using Early Posterior Negativity (EPN) amplitudes in silent reading task and found that individuals with functional gastrointestinal disorders (FGID) exhibited elevated EPN across various types of words, including symptom-related, emotionally neutral, and emotionally negative words. This general increase in EPN suggests a non-specific hyper-reactivity to stimuli at a pre-attentive level in FGID sufferers, indicative of broad, heightened unconscious attentional processing. Additionally, EPN amplitude correlated with psychosocial factors, where higher EPN levels were associated with lower health anxiety and coping through planning. These findings imply that the pre-attentional bias in IBS may also relate to coping styles, as those with high EPN seem less engaged in planning-based coping, reflecting a complex interaction between heightened unconscious attention and psychosocial factors. Together, these studies reveal that IBS patients display an exaggerated pre-attentional response to both sensory and GI-specific cues, with implications for symptom persistence and heightened symptom perception. The heightened sensitivity of pre-attentional systems likely contributes to an ongoing hypervigilance, where symptom-related stimuli are prioritized unconsciously, which can exacerbate symptom awareness and anxiety. Furthermore, the observed sex differences and correlations with psychosocial factors highlight that pre-attentional processing in IBS may not only be heightened but also modulated by individual factors, making it a nuanced component of the disorder’s symptomatology.

### 3.4. Neurofunctional Evidences in Attention Domain in IBS

A total of eight studies investigated neural mechanisms in IBS patients, using EEG/ERP, fMRI, and PET methodologies to explore sensory processing, attention in IBS patients. Five studies employed EEG/ERP methods: Blomhoff et al. (2001) and Berman et al. (2002) focused on sensory gating and attentional responses to stimuli Rustamov et al. (2020) examined pain-related brain activity modulation, while Ejova et al. (2021) and Sharma et al. (2021) assessed attentional biases and cognitive processing in IBS patients. Three fMRI studies were conducted: Andresen et al. (2005) explored sensory and emotional processing, Hubbard et al. (2015) assessed attention-related brain network activity, and Hong et al. (2016) investigated responses to contextual threats and pain anticipation. Finally, only Naliboff et al. (2008) utilized PET to examine changes in brain regions involved in pain processing and hypervigilance over time in IBS patients.

#### 3.4.1. The Electroencephalogram (EEG) Studies

Studies by Blomhoff et al. (2001) and Berman et al. (2002) show that IBS patients have increased responses to both GI-specific and general sensory stimuli. Specifically, enhanced N1 and P1 amplitudes suggest that IBS patients experience a augmented preattentive central nervous system response, indicating a potential deficit in sensory gating and habituation to threat stimuli. Blomhoff et al. further suggest that the IBS-PA (IBS with Phobic Anxiety) subgroup exhibited elevated N1 amplitudes across stimulus types, Showing enhanced attentional response in the context of anxiety. This responsiveness is possibly influenced by increased visceral sensitivity and anxiety-related modulation of the gut-brain axis.

Both Blomhoff et al. (2001) and Rustamov et al. (2020) showed that IBS patients have difficulties with selective attention and sensory modulation. For instance, Rustamov et al. reported that IBS patients exhibited a deficit in pain inhibition mechanisms, with reduced modulation of pain-related ERPs like N100, P260, and gamma oscillations, indicating a lack of effective pain gating. This supports the observation of increased gamma activity in pain-processing areas, which reflects an impaired capacity to downregulate pain sensitivity in IBS patients. Ejova et al. (2021) further highlighted a conscious and unconscious attentional bias toward symptom-related and emotionally negative cues in IBS patients, witnessed in increased occipital Early Posterior Negativity (EPN) and decreased N400 amplitudes for emotionally negative stimuli.

#### 3.4.2. Functional Magnetic Resonance Imaging (fMRI) Studies

In fMRI studies, Hubbard et al. (2015) observed altered brain activation in the anterior midcingulate cortex (aMCC) and insula during tasks requiring sustained attention, with IBS patients exhibiting greater activation in these regions even without significant changes in task performance. This heightened activation could signify that maintaining sustained attention in IBS requires additional cognitive resources due to ongoing interoceptive processing demands. The inability to fully disengage from internal sensations may create a cognitive load that diminishes efficiency in sustained attentional tasks.

Naliboff et al. (2006). Hong et al. found that IBS patients exhibited increased activation in the amygdala, insula, and other brain regions associated with threat and salience when faced with potential pain or discomfort. This increased brain activity reflects a tendency to anticipate and overestimate potential pain, which contributes to hypervigilance and increased attention to bodily sensations. Similarly, Naliboff et al. observed persistent activation in pain-processing areas such as the anterior insula and thalamus, but with a reduced response in limbic areas involved in vigilance over time, suggesting that chronic hypervigilance in IBS possibly is not diminished easily, even when individuals with IBS is exposed repeatedly to visceral stimuli.

#### 3.4.3. PET Study

The only PET study found in this systematic review was done by Naliboff et al. (2006) conducting a PET imaging over a 12-month period to analyze activation in pain-processing and vigilance-related circuits in response to rectal distention in IBS patients. The study showed persistent activity in areas such as the anterior insula and thalamus during rectal distension while the imaging sessions was taking place. These regions are involved in sensory aspects of pain processing and may be relevant for the orienting domain of attention, where sensory information is directed toward relevant bodily sensations. There was a significant decrease in activation in areas associated with vigilance, arousal, and emotional responses, particularly the amygdala and dorsal anterior cingulate cortex (dACC). This reduction in activity was observed during both anticipation and experience of rectal distension. This reduction in activation in vigilance-related areas was observed during both actual and anticipated visceral distention. However, over time, patients seemed to show less hypervigilance to expected and experienced abdominal discomfort, possibly indicating a form of habituation to the stimulus.

## 4. Discussion

The aim of present study is to review the literature on cognitive attention and its aspect in IBS patients. 24 relevant studies were identified, which employed a total of six different measures. 10 studies used neuroimaging measures or a combination of neuroimaging and behavioral measures of attentional (Andresen et al., 2005; Berman et al., 2002; Blomhoff et al., 2001; Ejova et al., 2021; Hong et al., 2016; Hubbard et al., 2015; Naliboff et al., 2008; Rustamov et al., 2020; Sharma et al., 2021), whereas the remaining nine used behavioral measures. five studies did not find a significant difference in aspects of attention between IBS and healthy groups (Attree et al., 2003; Berrill et al., 2013; Farup & Hestad, 2015; Kennedy et al., 2014; Sharma et al., 2021).

The finding of present systematic review shows an interplay of attentional processes in IBS emphasizing interactions with brain-gut axis dysregulation. Regarding the results of reviewed studies, while IBS patients did not have broad cognitive attention deficits, the demonstrated often attention biases, hypervigilance and pre-attentional processing disruptions, especially if symptom-related and negative cues were present, which may offer insight into cognitive mechanisms may contribute to symptom aggravation and durability of irritable bowel syndrome, also the emotional and psychological difficulties that associated with this IBS.

The existing literature show that general attention is not significantly impaired in IBS patients (Billing et al., 2024), and that the observed differences are more likely to be related to secondary effects of anxiety and depression(Farup & Hestad, 2015). In contrast, IBD patients show greater deficits in cognitive processing and general attention, which are characteristic of organic diseases (Sharma et al., 2021).

Selective attention in IBS is specifically focused towards symptom-related stimuli, leading to heightened symptom perception, anxiety, and hypervigilance (Gibbs-Gallagher et al., 2001; Kennedy et al., 2014; Thijssen et al., 2010). Depression, anxiety, and visceral sensitivity moreover modulate these biases, particularly in response to threat-related cues differently from that observed in healthy controls as seen in Tkalcic et al. (2014) and Elsenbruch et al. (2010) Additionally, IBS patients also display attention biases toward emotionally negative stimuli, marked by heightened neural responses and false-positive errors(Andresen et al., 2005; Phillips et al., 2014).

Emerging evidence suggests sustained attention deficits in IBS, as reflected in reaction time variability and poorer task performance (Berrill et al., 2013; Wong et al., 2019). Hypervigilance is a core feature of IBS, amplifying sensitivity to GI and emotional stimuli, driven by brain-gut interactions, but can be mitigated through habituation or distraction (Naliboff et al., 2008; Posserud et al., 2009).Moreover, IBS patients exhibit heightened pre-attentive processing, with biases toward symptom-related cues (Afzal et al., 2006) and increased sensitivity to sensory input, including elevated P1 and EPN amplitudes (Berman et al., 2002; Ejova et al., 2021). These findings suggest reduced sensory gating, broad hyper-reactivity, and sex-specific variations in prepulse inhibition (Kilpatrick et al., 2010).

Recent studies suggest that IBS patients exhibit an early attentional bias, as indicated by increased P1 and N1 amplitudes, leading to reduced sensory gating and increased preconscious awareness (Berman et al., 2002; Blomhoff et al., 2001). This is consistent with findings by Rustamov et al. (2020) and Ejova et al. (2021), who highlight the struggle with inhibitory control and attentional modulation in IBS patients, leading to a persistent focus on symptoms.

Neuroimaging studies, including those byAndresen et al. (2005), Hong et al. (2016), and Hubbard et al. (2015), have shown increased activity in emotional and sensory regions of the brain, indicating increased threat sensitivity and emotional processing of symptom-related information. This emotional bias enhances symptom perception and pain-related anxiety. Interestingly, Naliboff et al. (2008) found through PET studies that IBS patients may habituate attention over time, reduce hypervigilance, and improve emotional regulation. This adaptation could be clinically beneficial for IBS patients by reducing emotional burden and improving symptom management.

To elaborate on this point, it can be stated that the lack of sensory gating reflects an impaired ability to filter out repetitive or irrelevant stimuli, which could potentially make IBS patients hyper-sensitive toward bodily sensations. The amplified pre-attentional response supports the theory that altered brain-gut signaling increases baseline neural responsiveness, particularly to sensory and interoceptive cues, even when these stimuli are irrelevant or non-threatening (Mayer & Tillisch, 2011). This can explain why IBS patients tend to focus more on internal, symptom-related stimuli, as the brain-gut axis continuously signals heightened vigilance toward visceral sensations. Recently, supporting the mentioned notion, Formica et al. (2022) and Carpinelli et al. (2023) indicates that patients with Irritable Bowel Syndrome (IBS) exhibit significant sensitivity to internal sensations, particularly in relation to gastrointestinal symptoms. Specifically, it was found that there is a significant correlation between nausea and disgust, suggesting that IBS patients may experience heightened sensitivity to these internal sensations.

The brain-gut axis is a critical component in understanding the pathophysiology of IBS and its associated cognitive impairments, particularly in the domain of attention. Research has shown that dysregulation within the brain-gut axis can lead to alterations in both gastrointestinal and cognitive functions. For instance, neural network alterations that contribute to visceral hypersensitivity, frontal executive dysfunction and stress-related hippocampal-mediated cognitive alterations (Wong et al., 2019). This hypervigilance is thought to stem from heightened sensitivity in the salience network, which includes brain regions such as the anterior insula and anterior mid-cingulate cortex (Hong et al., 2016; Liu et al., 2016; Naliboff et al., 2008). These areas are involved in processing emotional and sensory information, suggesting that IBS patients may be more attuned to bodily sensations and potential threats, which can exacerbate their symptoms. Furthermore, the gut microbiota disruptions can lead to cognitive and behavioral changes, including attentional biases and executive dysfunctions (Quigley, 2018; Tang et al., 2023). This bias can lead to increased symptom perception and emotional distress, creating a cycle that reinforces the experience of pain and discomfort as shown in studies, such as Billing et al. (2024); Hubbard et al. (2015).

Neuroimaging studies further elucidate the relationship between the brain-gut axis and attention in IBS. For instance, IBS patients show disrupted functional connectivity density (FCD) in several brain regions, including decreased connectivity in the anterior midcingulate cortices and increased connectivity in sensorimotor cortices (Weng et al., 2017) .Additionally, studies integrating neuroimaging and fecal metabolite data have identified specific amino acid metabolites that correlate with IBS-specific brain connectivity changes. These metabolites may influence brain function directly or through peripheral mechanisms, supporting the brain-gut-microbiome model of IBS (Osadchiy et al., 2020).

### 4.1. Implications for the Treatment

Considering the results of the present systematic review interventions aimed at retraining attention could be beneficial. These might include exercises that help patients focus less on gastrointestinal symptoms and more on neutral or positive stimuli, thereby reducing the confirmatory bias towards negative information. Recently Tayama et al. (2017) suggest that Attention Bias Modification (ABM) could be a promising intervention for normalizing brain activity associated with attention and anxiety in IBS patients. Specifically, the observed changes in EEG patterns—namely, the increase in relative alpha power and decrease in beta power— indicate that ABM may help regulate neural processes tied to attentional focus and anxiety responses in IBS.

Given ABM’s focus on retraining attention patterns, it could be effectively combined with mindfulness or cognitive behavioral therapy (CBT) techniques. Such integration may provide IBS patients with a comprehensive toolkit for managing symptoms by not only adjusting their physiological responses but also modifying underlying thought patterns and attentional habits that reinforce anxiety and hypervigilance. Considering multiple studies have demonstrated that CBT significantly improves gastrointestinal symptoms in, CBT have the largest evidence base for IBS patients (Black et al., 2020).

Moreover, manipulating the gut microbiota with probiotics and prebiotics has emerged as a novel strategy to treat IBS. These interventions can help restore gut microbiota balance, potentially alleviating both gastrointestinal and cognitive symptoms (Pusceddu et al., 2018).

Considering the individual differences in symptom severity and cognitive processing, treatments should be personalized. For instance, women may require different strategies during different phases of their menstrual cycle due to hormonal influences on symptom perception (Kilpatrick et al., 2010). A novel intervention is personalized interventions and management approaches that significantly improves IBS-related symptoms and targeting individual symptom reduction in disorders of the IBS with different severity (Karakan et al., 2022; Schnedl et al., 2023).

### 4.2. Limitations

This systematic review on attention in IBS is subject to several limitations that should be considered when interpreting its findings. One major limitation is the heterogeneity of the included studies, which varied significantly in methodologies, patient populations, outcome measures, and cognitive assessment tools. Such variability complicates the synthesis of results and limits the ability to draw definitive conclusions. Additionally, the review includes a relatively small number of studies, which may not fully capture the breadth of existing literature on attention in IBS. This limited sample size restricts the robustness of the findings and may leave some aspects of attention in IBS underexplored.

Potential publication bias is also a concern, as studies reporting significant or positive findings are more likely to be published. This bias could skew the understanding of attention in IBS and overrepresent certain outcomes. Furthermore, the review focused exclusively on English-language articles, potentially excluding relevant studies published in other languages. This language limitation may narrow the breadth of the findings and exclude perspectives from diverse research contexts.

Another limitation is the predominance of cross-sectional designs among the included studies, which restricts the ability to infer causality between attention and IBS symptoms. Without longitudinal data, it remains challenging to determine whether attention biases are a cause or consequence of IBS symptoms. Moreover, the quality of the included studies varied, with some studies presenting methodological limitations that may affect the reliability and generalizability of their findings. This variability in study quality could influence the overall conclusions drawn from the review.

The review also focused primarily on attention-related cognitive domains, potentially overlooking other cognitive functions such as memory and executive function that may also play a role in IBS. This narrow focus limits the scope of the review and may fail to capture the full cognitive profile associated with IBS. Finally, the absence of longitudinal studies limits insights into how attention and biases evolve over time in relation to IBS symptoms and treatment, a perspective essential for understanding the dynamic nature of these cognitive processes.

These limitations underscore the need for further research to address these gaps, ideally through studies with more rigorous methodologies, larger sample sizes, and a broader exploration of cognitive functions. Addressing these limitations will enhance our understanding of attention in IBS and its implications for treatment.

### 4.3. Recommendations for Future Research

To advance our understanding of attentional biases in irritable bowel syndrome (IBS) and improve treatment strategies, several key areas warrant further investigation.

There remains a lack of comprehensive understanding regarding the specific cognitive mechanisms that contribute to attention in IBS. Future studies should focus on identifying and characterizing these mechanisms and investigating their impact on symptom perception and quality of life. Research should also explore how cognitive mechanisms underlying attentional biases interact with emotional and physiological factors, such as anxiety and visceral sensitivity, which are commonly observed in IBS patients.

Significant methodological variability across existing studies—such as differences in study populations, assessment tools, and outcome measures—further complicates efforts to draw consistent conclusions or synthesize findings across studies. To facilitate comparison and enable meta-analyses, it is crucial for future research to standardize methodologies, including consistent use of validated outcome measures and uniform protocols across studies. Such standardization will support more robust conclusions and offer greater insight into the cognitive aspects of IBS.

Longitudinal research is particularly needed to track cognitive changes over time in IBS patients, as well as their relationship to symptom severity and treatment outcomes. By following attention and biases longitudinally, researchers can better understand the stability of these features, observe how they interact with the natural progression of IBS, and evaluate the effectiveness of various treatments over time.

Additionally, there is a need for studies that integrate psychological, physiological, and neurocognitive data to more fully capture the multifaceted nature of IBS. Such interdisciplinary approaches could illuminate how cognitive processes relate to the brain-gut axis and contribute to IBS symptomology. Integrating these factors will also help clarify the reciprocal relationships between cognitive, emotional, and physical dimensions of IBS, potentially revealing new pathways for intervention.

Finally, the development and evaluation of targeted cognitive interventions, such as attention retraining and mindfulness practices, represent promising avenues for future research. By testing these approaches, researchers can assess their impact on modifying attentional biases and improving IBS outcomes. Addressing these research gaps will advance the understanding of IBS’s complex cognitive and neural underpinnings and pave the way for more personalized and effective therapeutic strategies.

## 5. Conclusion

This systematic review underscores the multifaceted nature of attention in patients with irritable bowel syndrome (IBS), revealing significant attentional biases alongside variations in other aspects of attention. The findings indicate that while IBS patients generally do not exhibit broad cognitive deficits, they demonstrate specific attentional biases, particularly towards negative emotional and gastrointestinal-related stimuli. This hypervigilance may exacerbate symptom perception and emotional distress, reflecting a complex interplay between cognitive processes and the brain-gut axis.

In addition to attentional biases, the review highlights the importance of sustained attention and selective attention in IBS. Furthermore, selective attention appears to be influenced by symptom-related cues, which may lead to difficulties in managing pain and other symptoms effectively.

In conclusion, the exploration of attentional mechanisms in IBS is crucial for developing a comprehensive understanding of the disorder. By integrating insights from cognitive psychology and neurobiology, future research can inform targeted interventions that address both the cognitive and emotional challenges faced by individuals with IBS. This holistic approach has the potential to enhance treatment outcomes and improve the quality of life for those affected by this complex condition.

